# Long non-coding RNA BCAR4 is required for efficient influenza A virus replication

**DOI:** 10.1101/2025.02.06.636885

**Authors:** Yi Cao, Alex W.H. Chin, Haogao Gu, Mengting Li, Leo L.M. Poon

## Abstract

Long non-coding RNAs (lncRNAs) regulate diverse biological processes, including influenza A virus (IAV) infection. However, the understanding of lncRNAs in IAV infection is limited. By using both bioinformatic analyses and virological assays, we showed that lncRNA *BCAR4* expression can be highly induced by infection of multiple different IAV subtypes. *BCAR4* was required for the propagation of IAV infection. Genetic inactivation of *BCAR4* inhibited IAV growth. Investigation of the IAV infection cycle revealed a suppressed IAV viral RNA transcription and replication, and attenuated viral protein synthesis in the *BCAR4*-deficient cells. *BCAR4* potentially interacted with cellular splicing-associated proteins and the activation of *BCAR4* was associated with influenza viral *NS* segment. These findings suggest the important role of lncRNA *BCAR4* in regulation of IAV infection.

**Importance:** Long non-coding RNAs (lncRNAs) serve as critical regulators in the biological processes of influenza A virus (IAV) infection. However, how lncRNAs engage IAV infection remains unclear. Here we show *BCAR4* as highly universally induced lncRNA in infection of multiple different IAV subtypes. Deletion of *BCAR4* reduced IAV multiplication. In the IAV infection cycle, *BCAR4* deficiency decreased IAV viral RNA transcription, replication and viral protein biosynthesis. *BCAR4* was potentially binding to the host RNA splicing associated protein, and the IAV viral NS segment is required for the activation of *BCAR4*. Our results highlight important regulation of *BCAR4* in IAV infection.

## Introduction

Influenza A virus (IAV) is a segmented, single strand RNA virus that can cause respiratory infection (1). The infection process of Influenza A Virus (IAV) involves a sequence of virological steps. These include the attachment and internalization of viral particles to the host cells, the viral RNA transcription and replication, viral protein synthesis, the assembly and virus budding of viral progeny (2). These virological activities, however, are either host machinery dependent or require cooperation from host factors (2–4). For instance, IAV infection exploits cellular RNA splicing machinery to promote viral RNA splicing (5, 6). Although hundreds of host factors required for efficient propagation of IAV have been identified by multiple screening or perturbation experimental systems (7–10), to date only a few of them have been well investigated.

Long non-coding RNAs (lncRNAs) regulate biological activities across diverse complex organisms (11, 12). It has broad functional engagement, ranging from regulation of genetic transcription in multiple levels to high dimensional organization of chromatin conformation (13). IAV infection can alter transcriptional profile of lncRNAs, which comprises an important part of the molecular signatures of host responses to the IAV infection (14). Because of the unusual profiling induced by infection, some lncRNAs have been investigated in detail, showing that different lncRNAs can act as either anti- or pro-viral infection factors, exemplified by *NEAT1* and *IPAN* (15, 16). However, the function of most lncRNAs in IAV infection remains unknown.

By exploring the transcriptomic profile of host lncRNAs in published datasets, we identified that lncRNA *BCAR4* is induced by infection of multiple different IAV subtypes. Deletion of *BCAR4* blocked IAV growth. In the IAV infection cycle, loss of *BCAR4* suppressed IAV viral RNA transcription and replication, as well as subsequent viral protein synthesis. *BCAR4* was potentially interacted with cellular RNA splicing-associated proteins and the activation of *BCAR4* was in association with IAV viral *NS* segment. This study revealed lncRNA *BCAR4* is required for IAV multiplication, highlighting the importance of lncRNA *BCAR4* in regulation of IAV infection.

## Results

### *BCAR4* is a highly universally triggered by infection of multiple different IAV subtypes

To explore universally differentially expressed host lncRNAs in infection of multiple different IAV subtypes, we previously analyzed publicly available high-throughput datasets that were produced from primary cells or cell lines of human origin infected with multiple different subtypes of IAV at multiply time points post-infection (17). By performing differential expression analysis in these datasets (summarized in Table 1), we determined that lncRNA *BCAR4* was up-regulated in infection by seven different IAV subtypes compared to mock infection (Fig. 1A). *BCAR4* showed an up-regulation trend over time during IAV infection and the expression fold change of *BCAR4* was significantly higher than mock infection around 6 to 12 hours-post infection (Fig. 1A). To validate this finding, we infected A549 or CALU-3 cells with either A/California/04/09 (H1N1) (CA04 hereafter) or A/Hong Kong/1/1968 (H3N2) (HK68 hereafter) strain with M.O.I. of 0.1 and examined the expression of *BCAR4* at multiple time points post-infection. Consistent with the induction of *BCAR4* in high-throughput datasets, the expression of *BCAR4* was remarkably higher in CA04 and HK68 infected cells compared to mock infected cells over time (Fig. 1B-C). We further performed a CA04 single cycle infection with high M.O.I. (M.O.I. of 5) in A549 cells to determine the earliest time points of *BCAR4* activation upon IAV infection. In line with previous results, the transcriptional activation of *BCAR4* started between 4 to 6 hours post-infection (Fig. 1D). Together, these results suggest that *BCAR4* is highly universally and reproducibly induced by infection of multiple different IAV subtypes.

**Fig 1.**
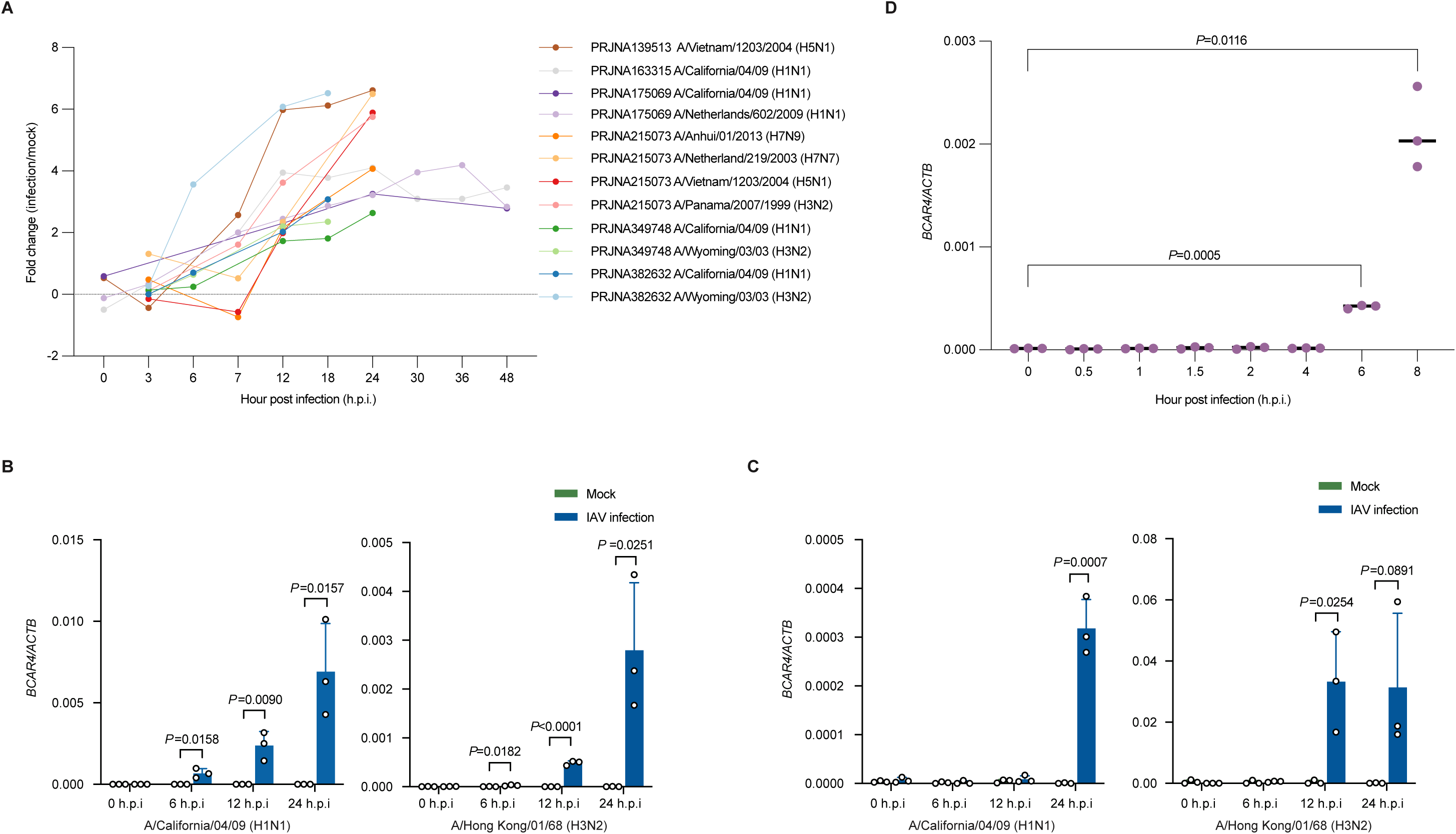
The up-regulation of *BCAR4* in infection of multiple different IAV subtypes. **(A)** The kinetics of fold change (infection/mock) in log2 of *BCAR4* expression in infection of multiple different IAV subtypes over time in included public datasets. The legend shows BioProject number and IAV subtype(s) used in the project. **(B)** The relative expression of *BCAR4* in either mock infected or A/California/04/09 (H1N1) (M.O.I. of 0.1) or A/Hong Kong/01/68 (H3N2) (M.O.I. of 0.1) infected A549 cells at 0, 6, 12 and 24 hours post-infection. The experiment was performed in triplicates. The statistical difference between two groups was calculated by using Student-t test. Two-sided significant *P*-value was reported in exact value. **(C)** The relative expression of *BCAR4* in either mock infected or A/California/04/09 (H1N1) (M.O.I. of 0.1) or A/Hong Kong/01/68 (H3N2) (M.O.I. of 0.1) infected CALU-3 cells at 0, 6, 12 and 24 hours post-infection. The experiment was performed in triplicates. The statistical difference between two groups was calculated by using Student-t test. Two-sided significant *P*-value was reported in exact value. (D) The relative expression of *BCAR4* in A/California/04/09 (H1N1) (M.O.I. of 5) infected A549 cells at 0, 0.5, 1, 1.5, 2, 4, 6, and 8 hours post-infection. The experiment was performed in triplicates. The statistical difference between two groups was calculated by using Student-t test. Two-sided significant *P*-value was reported in exact value.

**Table 1.** Studied influenza virus infection datasets.

| **Bioproject no.** | **Platform** | **Host** | **M.O.I.** | **Influenza virus subtype** |
| --- | --- | --- | --- | --- |
| PRJNA349748 | RNA-seq | HTBE | 5 | A/California/04/09 (H1N1) (3, 6, 12, 18, 24 h.p.i.),<br>A/Wyoming/03/03 (H3N2) (3, 6, 12, 18 h.p.i.) |
| PRJNA382632 | RNA-seq | MDM | 2 | A/California/04/09 (H1N1) (3, 6, 12, 18 h.p.i.),<br>A/Wyoming/03/03 (H3N2) (3, 6, 12, 18 h.p.i.) |
| PRJNA139513 | Microarray | CALU-3 | 1 | A/Vietnam/1203/2004 (H5N1) (0, 3, 7, 12, 18, 24 h.p.i.) |
| PRJNA163315 | Microarray | CALU-3 | 3 | A/California/04/09 (H1N1) (0, 3, 7, 12, 18, 24, 30, 36, 48 h.p.i.) |
| PRJNA175069 | Microarray | CALU-3 | 3 | A/Netherlands/602/2009 (H1N1) (0, 7, 12, 18, 24, 30, 36, 48 h.p.i.),<br>A/California/04/09 (H1N1) (0, 12, 24, 48 h.p.i.) |
| PRJNA215073 | Microarray | CALU-3 | 1 | A/Anhui/01/2013 (H7N9) (3, 7, 12, 24 h.p.i.),<br>A/Netherland/219/2003 (H7N7) (3, 7, 12, 24 h.p.i.),<br>A/Vietnam/1203/2004 (H5N1) (3, 7, 12, 24 h.p.i.),<br>A/Panama/2007/1999 (H3N2) (3, 7, 12, 24 h.p.i.) |

### Identification the dominant transcript of *BCAR4* by IAV infection

*BCAR4* is a human intergenic lncRNA at the short arm p13.13 of chromosome 16 with 5 potential transcripts encoded from 4 different exon regions of the genome based on reference annotation (Fig. S1A-B). The transcription start site (TSS), transcription termination site (TTS) and the transcript variants of *BCAR4*, however, are possibly imprecise, as lncRNAs are relatively not as well annotated as protein-coding genes (18). To identify the TSS of *BCAR4*, we performed 5’ Rapid Amplification of cDNA Ends (RACE) in CA04 infected or mock infected A549 cells. The PCR product of 5’ RACE was sequenced by Nanopore long reads sequencing (Fig. S2A). Most mapped reads were at 121 bp upstream and some reads mapped at 321 bp upstream in CA04 infected A549 cells, longer than previous annotated TSS (Fig. S3A). Fitting with previous outcomes, transcriptional profile of TSS of *BCAR4* was highly induced in IAV infection in comparison with mock infection (Fig. S3A). As a signal sequence of polyadenylation (AAUAAA) was detected in the 3’ end of *BCAR4*, we performed poly(A) tail-based 3’ RACE in A549 cells treated with either PBS, or LNA transfection, or CA04 infection (Fig. S3B). We originally planned to test three different LNA molecules specific for *BCAR4* to suppress its expression. However, except for LNA BCAR4#3, which surprisingly increased BCAR4 expression following transfection, the other LNA sequences had no noticeable impact on BCAR4 expression. As LNA *BCAR4*#3 transfection and CA04 infection both induced *BCAR4* transcription, we used both to increase the expression of *BCAR4* for 3’ RACE PCR. Consistent with reference annotation, a poly(A) tail at 3’ end was confirmed by Sanger sequencing (Fig. S3B).

As the *BCAR4* genome can encode many different RNA transcripts, we asked which transcript variants of *BCAR4* are dominant in IAV infection. To address the question, we constructed a bacterial clone library inserted with DNA amplicons of the consensus region between the first and last exon of *BCAR4* that was amplified with cDNAs reversely transcribed from mock infected or CA04 infected cells (Fig. S2C and Fig. S5). We sequenced 72 different plasmids from randomly picked bacterial clones in the library of mock or CA04 infection, and the count and the frequency of each *BCAR4* transcript variant was determined (Fig. 2). The distribution of counts across detected *BCAR4* transcript variants showed that transcript variant 5 was most dominantly expressed during IAV infection, followed by transcript variant 4, 6, 3 and 10 of *BCAR4* (Fig. 2). Notably, the transcript variant 5 of *BCAR4* was also the dominantly expressed RNA transcript in A549 cells under the state of non-infection (Fig. 2). Moreover, except for these reference transcripts, many novel transcript variants of *BCAR4* were also discovered, such as transcript variant 12 and 13, and there were transcripts that were similar to already annotated transcripts but featured minor variations in some regions (Fig. 2). To validate this part of data, we also performed Nanopore long reads sequencing in PCR products from a separated, above-described head-to-tail exon-exon junction PCR reactions. In accordance with previous outcomes, those transcripts of *BCAR4* were also detected (Fig. S6).

**Fig 2.**
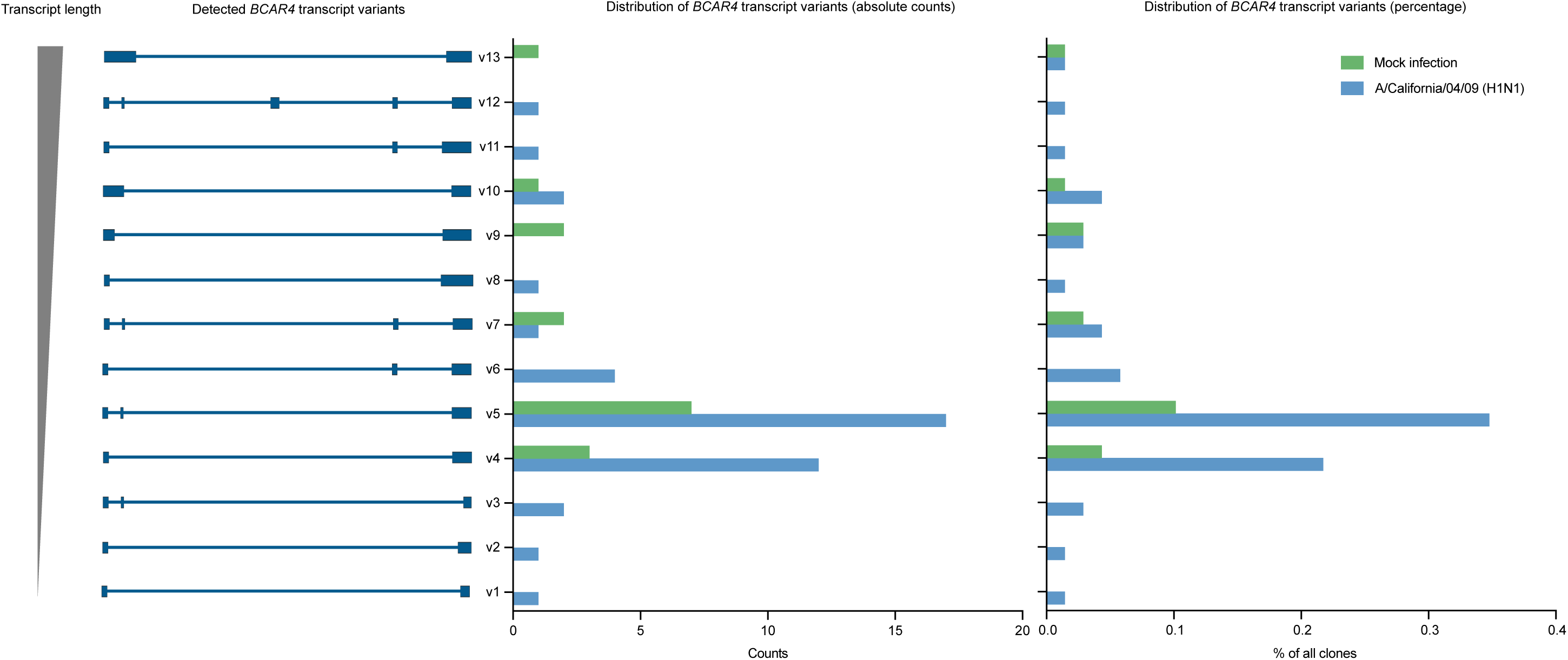
The distribution of *BCAR4* transcript variants. Identified transcript variants of *BCAR4* (the left) from clone library and the distribution of counts (the middle) and the percentage (the right) of each identified transcript in A/California/04/09 (H1N1) of M.O.I. of 1 infected or mock treated cells at 24 hours post-infection.

### Deletion of *BCAR4* suppresses influenza virus growth

Having observed that *BCAR4* was highly universally induced by IAV infection, we further asked whether *BCAR4* can affect IAV multiplication. Due to the capacity of lncRNAs to form secondary structures, the importance of different RNA regions of a lncRNA might vary, based on the spatially organized functional domains. Considering this, we co-transfected A549 cells with two pairs of CRISPR-CAS9 plasmids targeting the upstream and downstream of each exon of *BCAR4* (Fig. S7A). This allows the generation of cells with different combinations of deletion in the genome of each *BCAR4* exon region. We picked cell clones with full deletion in each exon of *BCAR4* in both alleles. These cell clones were marked as *BCAR4* exon1^F-/-^, *BCAR4* exon2^F-/-^, *BCAR4* exon3^F-/-^ and *BCAR4* exon4^F-/-^. The co-transfection also generated cell clones with full exon deletion in only one allele (single knock-out), including *BCAR4* exon1^+/-^, *BCAR4* exon2^+/-^ and *BCAR4* exon4^+/-^, or short length deletion in both alleles of targeted *BCAR4* exon regions, such as *BCAR4* exon3^S-/-^ and *BCAR4* exon4^S-/-^ cell clones. To gain a sense about the importance of these different lncRNA variants (Fig. S7B), we also explored these cells in following IAV infection experiments.

To examine if *BCAR4* affect the growth of IAV, we infected *BCAR4* exon1^F-/-^, *BCAR4* exon2^F-/-^, *BCAR4* exon3^F-/-^ and *BCAR4* exon4^F-/-^ cells and other cells carrying different combinations of *BCAR4* exon deletion with CA04 of M.O.I. of 1. There was an overall viral titer reduction in the supernatant of CA04 infected *BCAR4* KO cells compared to CA04 infected WT cells (Fig. 3A). Notably, except for the *BCAR4* exon1 KO pair, cells with homozygous full deletion in *BCAR4* exon, generated lower viral titer compared to their corresponding heterozygous *BCAR4* exon KO cells or short length homozygous *BCAR4* exon deletion cells, suggesting biallelic genetic inactivation of *BCAR4* in large length is required to produce inhibitory effect on IAV replication (Fig. 3A).

**Fig 3.**
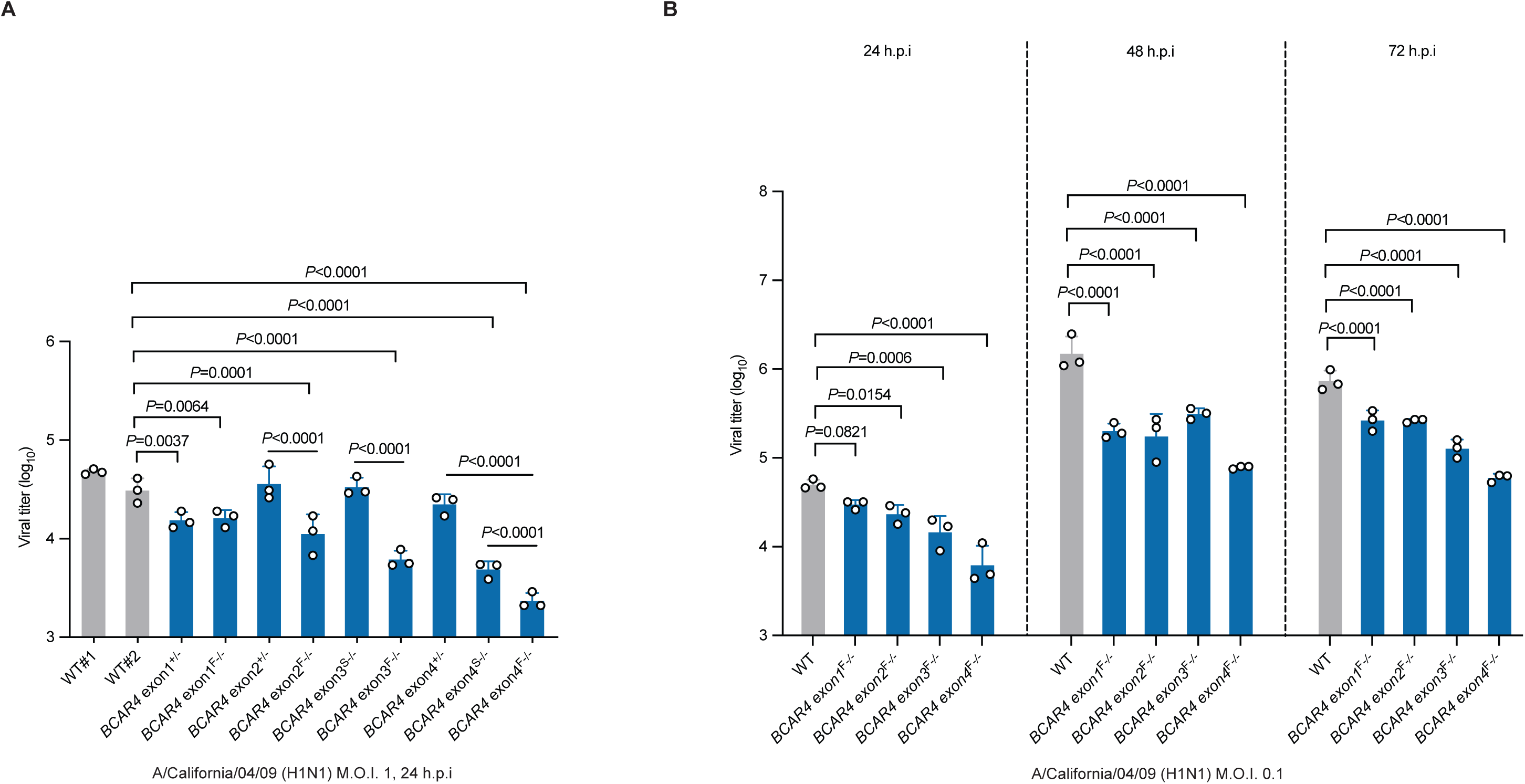
Loss of *BCAR4* suppresses IAV replication. **(A)** Viral titer in the supernatant of A/California/04/09 (H1N1) of M.O.I. of 1 infected A549 WT cells or infected A549 cells with different combinations of deletion in each exon of *BCAR4* at 24 hours post-infection. **(B)** Viral titer in the supernatant of A/California/04/09 (H1N1) of M.O.I. of 0.1 infected A549 WT cells or infected A549 cells with homozygous full deletion of each exon of *BCAR4* at 24, 48 and 72 hours post-infection. *BCAR4* exon1^+/-^, A549 cells with full deletion of *BCAR4* exon 1 in a single allele. *BCAR4* exon1^F-/-^, A549 cells with full deletion of *BCAR4* exon 1 in both alleles. *BCAR4* exon2^+/-^, A549 cells with full deletion of *BCAR4* exon 2 in a single allele. *BCAR4* exon2^F-/-^, A549 cells with full deletion of *BCAR4* exon 2 in both alleles. *BCAR4* exon3^S-/-^, A549 cells with short length deletion in *BCAR4* exon 3 in both alleles. *BCAR4* exon3^F-/-^, A549 cells with full deletion of *BCAR4* exon 3 in both alleles. *BCAR4* exon4^+/-^, A549 cells with full deletion of *BCAR4* exon 4 in a single allele. *BCAR4* exon4^S-/-^, A549 cells with short length deletion in *BCAR4* exon 4 in both alleles. *BCAR4* exon4^F-/-^, A549 cells with full deletion of *BCAR4* exon 4 in both alleles. Experiment was performed in triplicates. The statistical difference among three groups was calculated by using one-way ANOVA. The *LSD* test was used as a host hoc test for comparison between two groups. Two-sided significant *P*-value was reported in exact value.

To determine the growth kinetics of IAV in A549 cells in the presence or absence of each exon of *BCAR4*, we infected WT cells and cells with full deletion of each *BCAR4* exon with CA04 of M.O.I. of 0.1. The supernatant was collected at 24, 48 and 72 hours post infection. Consistent with the previous observations, reduced viral titer was detected in the supernatant of CA04 infected *BCAR4* exon1^F-/-^, *BCAR4* exon2^F-/-^, *BCAR4* exon3^F-/-^ and *BCAR4* exon4^F-/-^ cells, compared to the viral titer in the supernatant of CA04 infected WT cells at each time point post-infection (Fig. 3B), suggesting each exon of *BCAR4* is functionally essential to IAV infection. Together, these data suggest the *BCAR4* is one of the important factors required for IAV propagation.

### Genetic inactivation of *BCAR4* inhibits viral RNA transcription, replication, and viral protein synthesis

As IAV replication was affected by the processes of the infection cycle, we ask if *BCAR4* can influence the infection cycle of IAV. We first investigated whether *BCAR4* can affect the early stage of infection cycle including IAV attachment and internalization. To investigate if *BCAR4* impacts the attachment of IAV viral particles, high M.O.I. (M.O.I. of 5) CA04 were incubated with WT cells or cells with excision of each *BCAR4* exon at 4 °C for one hour. Low temperature blocks the viral internalization but not the attachment of IAV to host cells. Compared to CA04 infected WT cells, most CA04 infected KO cells, including *BCAR4* exon1^F-/-^, *BCAR4* exon2^F-/-^, *BCAR4* exon3^F-/-^ and *BCAR4* exon4^S-/-^, showed comparable viral attachment, except for *BCAR4* exon4^F-/-^, which had higher viral attachment than infected WT cells (Fig. 4A). We next investigated the effect of *BCAR4* to IAV internalization. High M.O.I. (M.O.I. of 5) of CA04 were used to infect WT cells or cells carrying each *BCAR4* exon excision at 37 °C for one hour. After incubation, uninternalized viruses were inactivated by acidified sodium chloride solution (pH 2) and RNA from infected cells were harvested half hour later. The expression of the viral *M* gene was used to estimate the amount of internalized viruses. No significant difference was found in internalized viral RNA in CA04 infected *BCAR4* exon1^F-/-^, *BCAR4* exon2^F-/-^ and *BCAR4* exon4^S-/-^ cells, whereas CA04 infected *BCAR4* exon3^F-/-^ and *BCAR4* exon4^F-/-^ had higher viral internalization of CA04, compared to CA04 infected WT cells (Fig. 4B).

**Fig 4.**
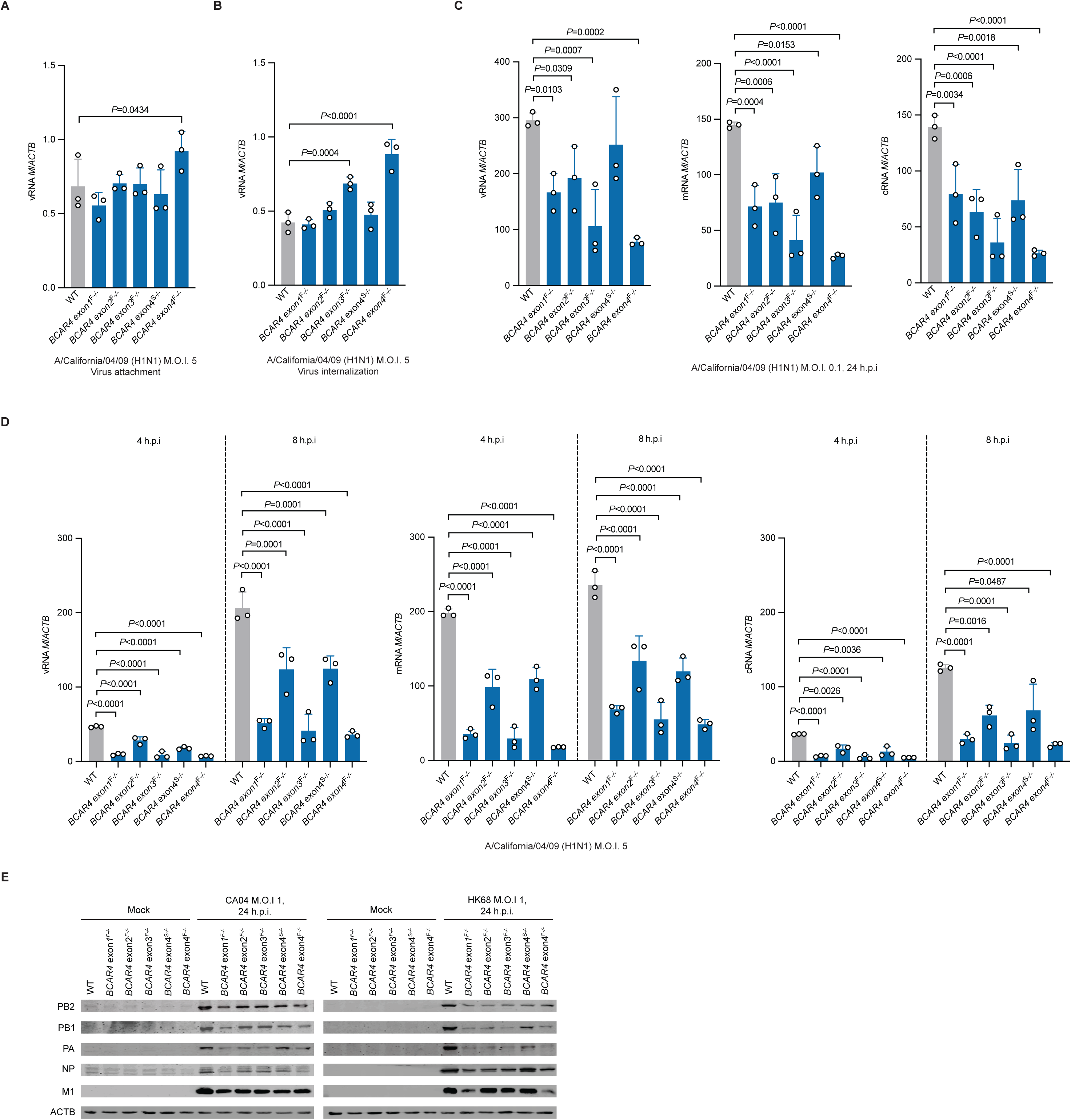
Lack of *BCAR4* inhibits IAV viral RNA transcription, replication, and viral protein synthesis. **(A)** The relative expression of viral *M* gene in attached viruses of A/California/04/09 (H1N1) (M.O.I. of 5) infected A549 WT cells or infected A549 cells with each *BCAR4* exon deletion at one-hour post-infection. **(B)** The relative expression of viral *M* gene in internalized viruses of A/California/04/09 (H1N1) (M.O.I. of 5) infected A549 WT cells or infected A549 cells of each *BCAR4* exon deletion with at 1.5 hours post-infection. **(C)** The relative expression of vRNA (the left), mRNA (the middle) and cRNA (the right) of viral *M* gene in A/California/04/09 (H1N1) (M.O.I. of 0.1) infected A549 WT cells and infected A549 cells with each *BCAR4* exon deletion at 24 hours post-infection. **(D)** The relative expression of vRNA (the left), mRNA (the middle) and cRNA (the right) of viral *M* gene in A/California/04/09 (H1N1) (M.O.I. of 5) infected A549 WT cells and infected A549 cells of each *BCAR4* exon deletion at 4 and 8 hours post-infection. **(E)** The expression of viral protein PB2, PB1, PA, NP and M1, as well as cellular protein ACTB in A/California/04/09 (H1N1) (M.O.I. of 1) or A/Hong Kong/1/1968 (H3N2) (M.O.I. of 1) infected A549 WT cells or infected A549 cells with each *BCAR4* exon deletion at 24 hours post-infection. *BCAR4* exon1^F-/-^, A549 cells with full deletion of *BCAR4* exon 1 in both alleles. *BCAR4* exon2^F-/-^, A549 cells with full deletion of *BCAR4* exon 2 in both alleles. *BCAR4* exon3^F-/-^, A549 cells with full deletion of *BCAR4* exon 3 in both alleles. *BCAR4* exon4^S-/-^, A549 cells with short length deletion in *BCAR4* exon 4 in both alleles. *BCAR4* exon4^F-/-^, A549 cells with full deletion of *BCAR4* exon 4 in both alleles. Gene expression experiments were performed in triplicates. The statistical difference among three groups was calculated by using one-way ANOVA. The *LSD* test was used as a host hoc test for comparison between two groups. Two-sided significant *P*-value was reported in exact value.

As *BCAR4* does not block virus entry and attachment, we continue to examine following stages of the infection cycle, including viral RNA transcription and replication. We infected WT cells and each *BCAR4* exon deficient cells with CA04 of M.O.I. of 0.1. and harvested cells at 24 hours post infection. Quantification of vRNA, mRNA or cRNA of CA04 *M* gene showed an overall declined viral RNA transcription and replication in CA04 infected *BCAR4* exon1^F-/-^, *BCAR4* exon2^F-/-^ or *BCAR4* exon3^F-/-^, *BCAR4* exon4^F-/-^ cells compared to infected WT cells (Fig. 4C). To determine if the loss of *BCAR4* impairs viral RNA transcription and replication in single cycle infection, we infected WT cells or cells with each *BCAR4* exon deletion with CA04 of M.O.I. of 5, and harvested infected cells at 4 and 8 hours post-infection. There was a total attenuated vRNA, cRNA and mRNA of viral *M* gene in CA04 infected cells with each *BCAR4* exon deletion in comparison with CA04 infected WT cells at 4 and 8 hours post-infection (Fig. 4D), suggesting that genetic inactivation of *BCAR4* can inhibit viral RNA transcription and replication in the single infection cycle. We also investigated the viral protein synthesis in CA04 or HK68 infected WT cells or infected cells with each *BCAR4* exon deletion. Consistent with the result of *BCAR4* deletion inhibiting viral RNA transcription and replication, the overall expression of multiple viral proteins, including PB2, PB1, PA, NP and M1, was remarkably lower in CA04 or HK68 infected cells carrying each *BCAR4* exon deletion compared to infected WT cells (Fig. 4E). Collectively, these data suggest that loss of *BCAR4* inhibits IAV viral RNA transcription and replication, as well as subsequent viral protein synthesis.

### *BCAR4* is potentially interacted with cellular splicing-associated proteins

To determine proteins that potentially interacted with *BCAR4*, we performed *BCAR4* RNA pulldown assay *in-vitro* and identified *BCAR4* interacted protein by Mass Spectrometry (MS). We *in-vitro* transcribed the sense RNA strand (functional RNA sequence) of the dominant transcript variant of *BCAR4* (transcript variant 5). The antisense RNA strand was reversely transcribed for experimental control and normalization. RNA probes of sense or antisense of the dominant *BCAR4* transcript were incubated in cell lysates from either mock infected or CA04 (M.O.I. of 1) infected A549 cells. After incubation, *BCAR4* RNA interacted proteins were identified by MS. To directly compare *BCAR4* RNA interacted proteins in IAV infection and non-infection state, we used the FC-FC plot to exhibit the fold change of enriched proteins in the *BCAR4* functional sense strand versus its antisense strand in each experimental condition (Fig. 5). RNA pulldown assay showed that proteins potentially interacted with the dominant transcript of *BCAR4* were mostly cellular RNA-splicing associated, such as hnRNPH1 (19, 20), hnRNPH3 (19), FUBP1 (21) and SNRPA (22), and these identified proteins were mostly consistent in both CA04 infection and mock infection (Fig. 5). Besides, no IAV viral protein was significantly enriched in the *BCAR4* probe. These data implies that *BCAR4* may interact with host cellular RNA splicing machinery related proteins.

**Fig 5.**
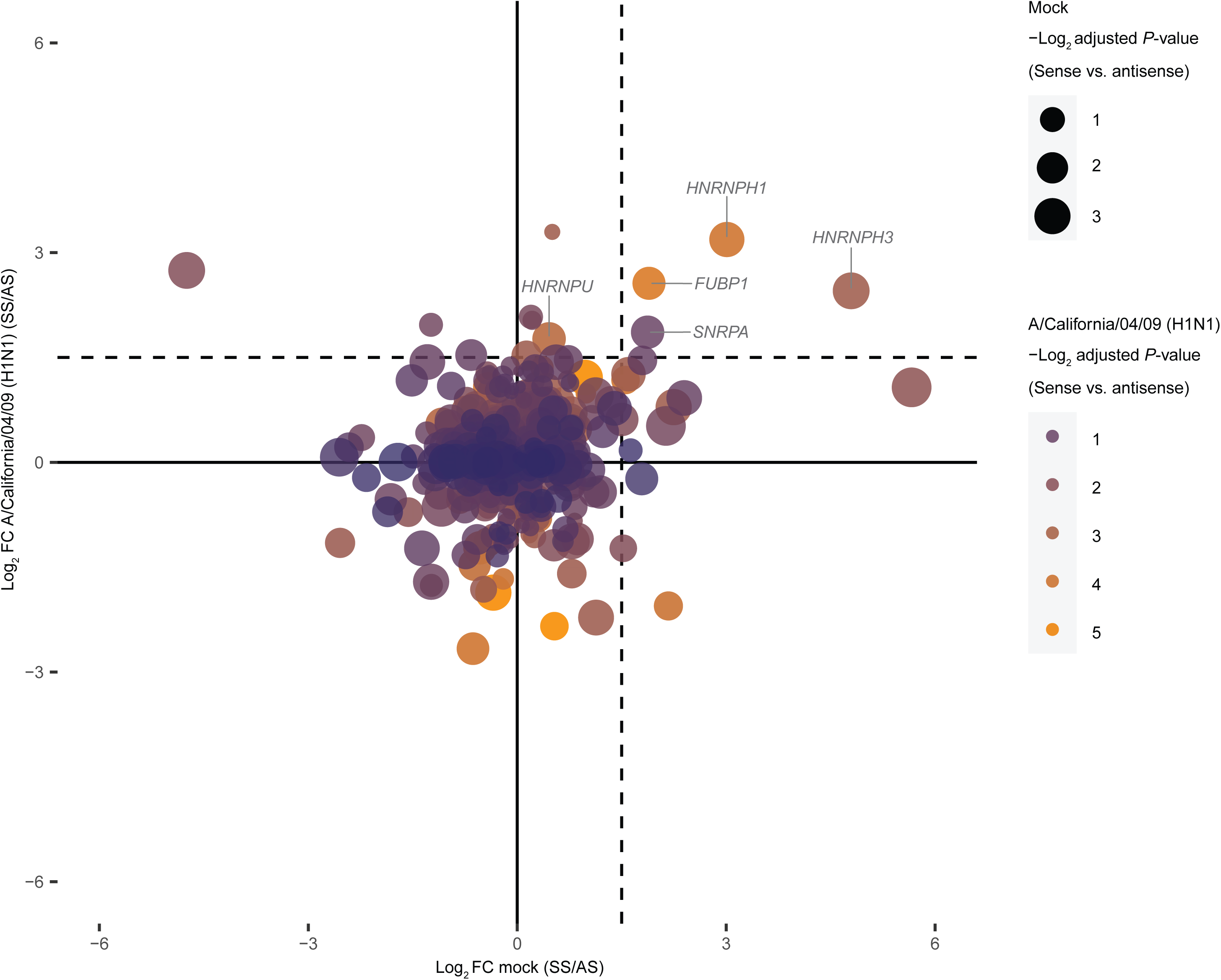
*BCAR4* is potentially interacted with cellular RNA-splicing associated proteins. FC-FC plot shows Mass Spectrometry (MS) identified proteins that enriched in dominant *BCAR4* functional sense RNA probes (*BCAR4* transcript variant 5) normalized with control antisense strand from cell lysates of PBS mock treatment (X axis) or A/California/04/09 (H1N1) (M.O.I. of 1) infection (Y axis) at 24 hours post-infection. The *P*-value in the mock group and infection group visualize by the size or the color of the dot respectively. The dashed line indicates a fold change value of 1.5. The RNA-pulldown MS was performed in triplicates.

### The activation of *BCAR4* is associated with influenza viral *NS* segment

*BCAR4* is activated by infection of multiple different IAV subtypes, but whether the activation of *BCAR4* is related to innate immune response is unknown. To this end, we treated A549 cells with multiple different immune stimuli individually, including interferon α, interferon β, interferon γ and LPS, with a gradient of different concentration. Compared to positive controls, such as *ISG15* and *MX1* that were dramatically induced by immune stimulation, the expression of *BCAR4* were mostly unchanged in response to the stimuli at 6 and 24 hours post-treatment (Fig. S8A and S8B). We also analyzed multiple published high-throughput datasets that generated from interferons or LPS stimulation experiments (Table 2). Unlike the remarkably up-regulated *BCAR4* identified in the infection datasets (Fig. 1A), the profile of *BCAR4* in these immune stimulation datasets were mostly unmeasurable (Fig. S8C), suggesting immune stimuli are not able to trigger the transcriptional activation of *BCAR4* from the low basal expression level. Considering that *BCAR4* might be responsive to other types of stimuli except for interferons, we also treated A549 cells with conditioned medium, a virus-free cell culture medium collected from IAV infected cells containing a pool of cellularly secreted antivirals, proinflammatory molecules and other infection-related factors. The expression of *BCAR4* was also not triggered by conditioned medium in comparison with remarkably triggered positive control genes including *ISG15* and *MX1* over time (Fig. S9). These results suggest that the activation of *BCAR4* is independent of immune responses.

**Table 2.** Studied RNA-seq datasets generated from hosts immune stimulation experiments.

| Bioproject no. | Platform | Host | Treatment |
| --- | --- | --- | --- |
| PRJNA788896 | RNA-seq | Sigmoid colon organoid | Interferon-β; Interferon-γ |
| PRJNA768480 | RNA-seq | A549 | Interferon-γ |
| PRJNA667475 | RNA-seq | A549 | Interferon-β |
| PRJNA795806 | RNA-seq | PBMC | LPS |

Given the clue that *BCAR4* may potentially interact with cellular RNA splicing associated proteins, we asked if *BCAR4* involves RNA splicing. To test the hypothesis, we treated A549 cells with Madrasin, which is a potent cellular spliceosomes inhibitor that can block the assembly of spliceosomes in the early stage (23). Interestingly, treating cells with Madrasin activated the transcription across multiple *BCAR4* genomic regions (Fig. 6A). However, we doubted if the up-regulation is *BCAR4* specific, as dysfunction of spliceosomes might cause insufficient production of mature RNA, potentially leading to cellular stress that may push cells to increase the global transcription for more functional RNA. We therefore examined the expression of some other types of RNA genes or protein-coding genes. We detected the expression of housekeeping gene *GAPDH*, and found no significant difference in expression of *GAPDH* between Madrasin treated cells compared to untreated cells (Fig. 6A). The expression of *USP30-AS1*, an interferon signaling dependent lncRNAs (17), was also not influenced by the spliceosome inhibition (Fig. 6A). We noticed that the expression of some small nuclear RNAs (snRNAs) including *RNU2* and *RNU6*, which are components of spliceosomes (24), were higher in Madrasin treated cells compared to the mock (Fig. 6A), suggesting that the transcriptional change of genes in response to Madrasin might more relate to RNA splicing processing.

**Fig 6.**
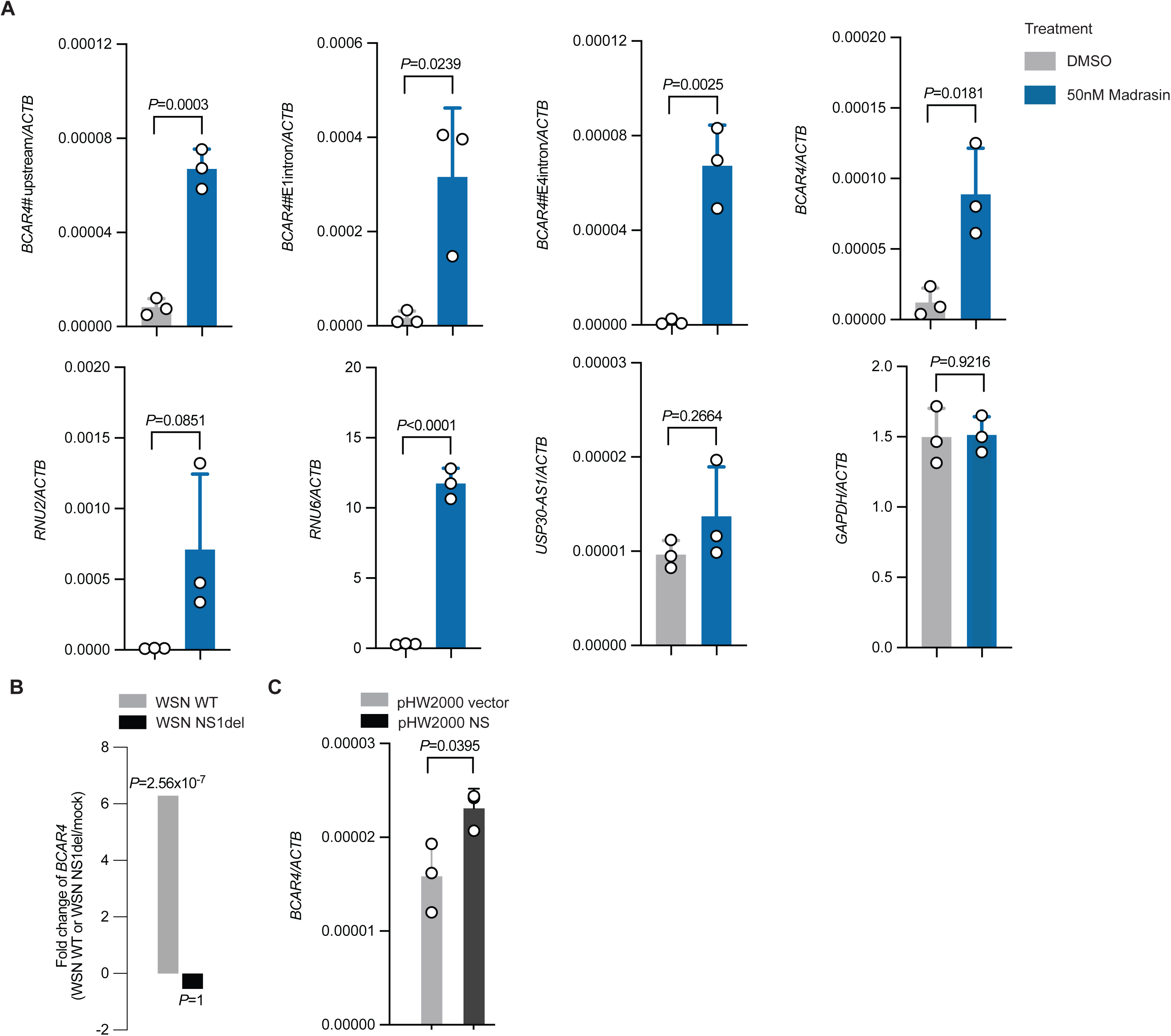
The activation of *BCAR4* is associated with the IAV *NS* segment. **(A)** The relative expression of *BCAR4* of multiple regions and housekeeping gene *GAPDH*, lncRNA *USP30-AS1* and snRNA *RNU2* and *RNU6* in 50 nM Madrasin treated A549 cells compared to DMSO treated A549 cells at 24 hours post-treatment. **(B)** Expression fold change (log2) of *BCAR4* in WSN WT infected cells versus mock infected cells and expression fold change (log2) of *BCAR4* in cells infected with NS1 segment deletion WSN strain compared to mock infected cells in a published RNA-seq dataset (BioProject PRJNA656848). **(C)** The relative expression of *BCAR4* in HEK 293T cells transfected with either empty pHW2000 plasmids or pHW2000 plasmids inserted with IAV viral *NS* segment. The experiment was performed in triplicates. The statistical difference between two groups was calculated by using Student-t test. Two-sided significant *P*-value was reported in exact value.

Considering that IAV viral NS1 protein also acts as a spliceosome antagonist, a similar role to Madrasin during infection, we hypothesized that the activation of *BCAR4* may be associated with viral NS segment. To test the hypothesis, we investigated a published dataset that produced from WT IAV WSN infected cells compared to cells infected with WSN with NS segment deletion (from BioProject PRJNA656848). Compared to WSN WT infection that triggered the up-regulation of *BCAR4* around 6-fold compared to mock infection, NS1-deficient WSN failed to trigger the activation of *BCAR4* in comparison with mock infection (Fig. 6B). To validate this result, we also transfected plasmids expressing IAV NS segment into HEK 293T cells and found transfection of viral NS segments enhanced the expression of *BCAR4* relative to the transfection of empty vector plasmids (Fig. 6C). Taken together, those results suggest that the activation of *BCAR4* is in association with IAV viral *NS* segment.

## Discussions

It is unclear how universally induced lncRNAs by different IAV subtypes engage IAV infection. By exploring multiple published datasets, we identified and validated lncRNA *BCAR4* that was highly universally up-regulated during infection of multiple different IAV subtypes. Due to this unique signature profile, we are interested in how *BCAR4* regulates IAV infection. As most lncRNAs are not as well annotated as protein-coding genes (18), it is necessary to identify the TSS, TTS and transcript variants of *BCAR4*. To address the issue, we performed 5’/3’ RACE for *BCAR4* together with either Sanger sequencing or Nanopore long reads sequencing. It was identified that *BCAR4* had a longer 5’ end than reference annotated and a poly(A) tail at its 3’ end. Through head-to-tail exon-exon junction PCR, we determined dominant transcript variants of *BCAR4* in IAV infection or non-infection. It was identified that transcript variant 5 of *BCAR4* was the most dominantly expressed RNA transcript in both IAV infection and non-infection condition, and many novel transcript variants of *BCAR4* were discovered for the first time.

To further investigate the phenotypic function of *BCAR4* in IAV infection, we generated A549 cells with full deletion in each *BCAR4* exon by using CRISPR-CAS9 gene editing. This allowed us to explore the importance of different RNA regions of *BCAR4* in IAV infection. To determine if *BCAR4* is required for IAV growth, we infected WT cells or cells of each *BCAR4* exon deletion with IAV, and found deficiency of any exon of *BCAR4* can suppress IAV multiplication. We further investigated whether *BCAR4* plays role in any specific viral events, which could account for the reduced virus replication. We first examined the capacity of attachment and internalization of IAV to host cells in the presence or absence of *BCAR4* and observed mostly comparable attached or internalized viruses in cells with each *BCAR4* exon deletion compared to WT cells. We then performed a multiple cycle IAV infection in WT cells and cells with deletion of each *BCAR4* exon to determine if *BCAR4* can influence viral RNA transcription and replication. Deletion of *BCAR4* inhibited viral RNA transcription and replication in infected *BCAR4* exon deficiency cells in multiple cycle infection in comparison with WT cells. Further investigation revealed that *BCAR4* also remarkably reduced viral RNA transcription and replication in high M.O.I. mediated IAV single cycle infection. We also examined the synthesis of viral proteins and found that there was an overall reduced viral protein production in IAV infected *BCAR4* exon deletion cells compared to infected WT. Taken together, these data suggest *BCAR4* is essential to IAV infection.

To gain more insight about proteins that potentially interacted with *BCAR4*, we performed *in-vitro* RNA pulldown assay using the dominant transcript variant of *BCAR4*, together with Mass Spectrometry for protein identification. Cellular RNA splicing associated proteins, such as spliceosome component proteins hnRNPH1/3 (19), but not IAV viral proteins, were enriched in *BCAR4* RNA probes in the state of IAV infection or non-infection. Because *BCAR4* is highly universally up-regulated by infection of multiple different IAV subtypes, we investigated which biological factors control the activation of *BCAR4*. To test if *BCAR4* involves immune responses, we examined the expression of *BCAR4* by different stimuli including interferon, LPS and conditioned medium. We found *BCAR4*, however, did not respond to these immune stimulations. Given the clue that *BCAR4* may interact with RNA-splicing associated cellular proteins, it is possible that *BCAR4* involves RNA splicing processes. To test this notion, we treated A549 cells with a potent spliceosome inhibitor Madrasin. Interestingly, *BCAR4* was significantly activated in cells treated with Madrasin, compared to DMSO treated cells. This activation of *BCAR4* is more related to spliceosome inhibition, as supported by activated expression of *RNU2* and *RNU6*, which are also core components of spliceosomes, but not genes that involve other biological actions, such as *USP30-AS1*, an interferon stimulated genes and *GAPDH*, a metabolic regulatory housekeeping gene. As inhibition of spliceosomes activates *BCAR4*, we turned our focus on IAV viral *NS* segment, as it serves a similar role as a spliceosome antagonist. We investigated a published datasets that generated from cells infected by either WSN WT or WSN with *NS1* segment deletion, and found compared to the remarkable induced expression of *BCAR4* in response to WSN WT infection, cells infected with WSN *NS1* segment deletion strain was not able to triggered the activation of *BCAR4*. Consistently, transfection of plasmids with IAV *NS* segment promoted the expression of *BCAR4* in comparison with cells transfected with empty vectors. These data suggests that the disruption of host RNA splicing machinery, either by pharmacological spliceosome inhibition or viral NS antagonism during IAV infection, is associated with the genetic activation of *BCAR4*, which later binds to spliceosomal components, such as hnRNPH1/3 and SNRPA (19, 20, 22). We have two non-mutually exclusive hypotheses to explain the link between the behavior of *BCAR4* and IAV replication. Firstly, as a factor engaging RNA splicing machinery, *BCAR4* has restricted physiological expression, suggesting a limited demand of *BCAR4* for canonical processing of RNA splicing. Inhibition of spliceosomes by IAV NS segment drives the production of *BCAR4* that might offer an alternative pathway for RNA splicing, a cellularly controlled non-canonical process but can help the facilitation of viral RNA processing during IAV infection. Second, some *BCAR4* interacted proteins, such as hnRNPH1, have antiviral activity in IAV infection (25, 26). Promoted by IAV, *BCAR4* might competitively bind to these antiviral proteins, attenuating their capacity to interact and inhibit viral components. However, further investigations are required to elucidate the potential mechanism of *BCAR4* in regulation of cellular and viral RNA splicing.

In conclusion, we identified a highly universally induced lncRNA *BCAR4* by infection of multiple different IAV subtypes, which is required for IAV multiplication, suggesting the important role of *BCAR4* engaging in IAV infection.

## Methods

### Viruses and cells

Influenza A viruses of A/California/04/01 (H1N1) and A/Hong Kong/1/1968 (H3N2) were rescued from the reverse genetic plasmids system as previously described (27). Virus was propagated in embryonated eggs. In brief, 1x10^3^ pfu/ml viruses in 100 μl PBS were first inoculated into ten-days embryonated eggs. Inoculated eggs were next incubated at 37 °C with a 55-60% humidity environment for two days. The eggs were then placed in -20 °C for one hour. After that, allantoic fluid was collected and centrifuged at 1000 g for 10 mins at 4 °C to remove debris. Supernatant was collected and aliquoted into 1.5 ml screw cap tubes and stored at -80 °C freezer for future use. The viral titer of the virus stock was determined by plaque assay.

A549, CALU-3, HEK-293T and MDCK cells were cultured in Minimum Essential Media (MEM) (11095080, Thermofisher), supplemented with 10% heat inactivated fetal bovine serum (FBS) (16000044, Thermofisher) and 1% penicillin and streptomycin (P/S) (15140148, Thermofisher). CALU-3 cells were maintained in Dulbecco’s Modified Eagle Medium (DMEM) (11965092, Thermofisher), supplemented with 10% FBS and 1% P/S. All cells were kept in incubators at 37 °C with 5% CO_2_.

### Plasmids and antibodies

Plasmids applied for reverse genetic transfection were made as previously described (17). Plasmids of pSpCas9(BB)-2A-Puro (PX459) V2.0 for gene editing were obtained from Addgene repository (62988, Addgene) (28).

Primary antibodies used in the study were PB2 (Sc17603, Santa Cruz Biotechnology) (1:1000), PB1 (NR31691, BEI) (1:1000), PA (PA5-31315) (1:1000), NP (ab128193, Abcam) (1:2000) and M1 (Ac57881, Santa Cruz Biotechnology) (1:2000), ACTB (Sc47778, Santa Cruz Biotechnology) (1:5000). Secondary antibodies used include Secondary antibodies including IRDye® 680RD Donkey anti-Goat IgG (H + L) (926-68074, LI-COR) (1:2000), IRDye® 680RD Donkey anti-Mouse IgG (H + L) (926-68072, LI-COR) (1:2000), IRDye® 680RD Donkey anti-Rabbit IgG (H + L) (926-68073, LI-COR) (1:2000).

### Transfection

For transfection of plasmids inserted with viral *NS* segment of A/Puerto Rico/8/1934 (H1N1) to HEK 293T cells, cells were seeded into 24-wells plates with 70% confluence one day before transfection. 500 ng of pHW2000 plasmids were diluted in Opti-MEM (31985070, Thermofisher) containing PLUS reagent (15338100, Thermofisher) and incubated with Opti-MEM diluted LTX Lipofectamine reagent (15338100, Thermofisher) for 5 mins. The plasmids-PLUS-LTX mix were next vortexed briefly and then added into cells. Transfection medium was kept in cells for eight hours before being replaced by fresh medium. Transfected cells were harvested 24 hours after transfection. For transfection of locked nucleic acid (LNA), 25 nm of LNA were used for each transfection with RNAiMAX (13778030, Thermofisher) based on the manufacturer’s instructions.

### Generation of knock-out cell lines

To generate A549 cells with each exon deletion of *BCAR4*, four individual sgRNA plasmids were pooled to co-transfection targeting upstream (two sgRNAs) and downstream (two sgRNAs) of each *BCAR4* exon. 500 ng (125 ng for each PX459 plasmids containing single sgRNA) of PX459 plasmids were diluted in Opti-MEM (31985070, Thermofisher) containing PLUS reagent and Lipofectamine LTX reagent. The plasmids-PLUS-LTX mix were sufficiently mixed and then incubated at room temperature for 5 mins. After that, an incubated plasmids-PLUS-LTX mix was added into cells. The transfection medium was kept in cells for 8 hours before being replaced by fresh culture medium. Transfected cells were selected by treating with 2.5 μg/ml Puromycin for two days. Survived cells were washed three times with BPS and serially diluted into 96-wells for potential single cell clone formation. Single cell clones were determined under microscopy and picked for PCR-based genotyping validation.

### IAV infection

For single cycle IAV infection, multiplicities of infection (M.O.I.) of 5 were used. Cells were first washed twice with PBS to remove serum and diluted viruses in PBS were incubated with cells at 37 °C for one hour. Unattached viruses were then removed, and serum-free culture medium was refilled. Cells were maintained for further downstream analysis. Typically, cells were harvested no more than 8 hours post infection in high M.O.I. single cycle infection experiment. For multiple cycle IAV infection, M.O.I. ranging from 0.1 to 1 was used based on experiment purposes. The virus incubation steps were mostly the same as single cycle IAV infection and cells were harvested based on the purposes of downstream analyses. Typically, for immunoblotting, IAV of M.O.I. of 0.1 was used and cells were harvested at 24 hours post infection. For determining virus growth kinetics, IAV of M.O.I. of 0.1 was used and cells were harvested at 24, 48 and 72 hours post infection. For detection of virus attachment, IAV of M.O.I. of 5 were diluted in ice-cold PBS and kept at 4 °C until use. Cells were washed with ice-cold PBS twice to remove serum and then viruses were incubated with cells at 4 °C for one hour. Low temperature can inhibit IAV entry but not attachment to the cells. After incubation, cells were washed three times with ice-cold PBS and harvested for downstream analysis. For detection of virus internalization, cells were incubated with IAV of M.O.I. of 5 at 37 °C for 1 hour. After incubation, cells were first washed once with pH 2 normal saline and washed with PBS. Washing cells with normal saline of pH 2 can collapse uninternalized viruses. After washing, cells continued to incubate for 30 mins, allowing IAV to release their viral RNA into cytoplasmic space but not to replicate yet. Cells then were harvested for downstream analysis. For multiple cycle infection, TPCK-trypsin (4370285-1KT, Sigma) was added to the serum-free cell culture medium with 1 μg/ml for MDCK, HEK-293T and CALU-3 cells and 0.5 μg/ml for A549 cells.

### Plaque assay

MDCK cells were seeded into 6-well plates and cultured to 100% confluence. IAV were serially diluted in PBS from 10^-1^ to 10^-6^. Cell culture medium in full confluent MDCK cells was aspirated and MDCK cells were washed twice with PBS. Diluted IAV were serially added to each well and MDCK cells were next incubated at 37 °C for 1 hour. After incubation, unattached viruses were removed and MDCK cells were washed once with PBS. 2 ml of culture medium containing 1% SeaKem LE Agarose (50004, Lonza) with 1 ug/ml TPCK-trypsin was added to cells when the mix was at a mild temperature. The MDCK cells were incubated at 37 °C, 55-60% humidity incubator for two days. After that, MDCK cells were fixed with 10% formaldehyde for 1 hour and MDCK cells were stained with 1% crystal violet for 30 mins. Last, the crystal violet was removed, and 6-well plates were washed and dried for plaque counting.

### RNA extraction, PCR and qRT-PCR

Cells were lysed and RNA were isolated and purified by using RNeasy Mini Kit (74104, Qiagen) based on manufacturer’s instruction. Super Script II Reverse Transcriptase (18964071, Thermofisher) was used for reverse transcription of first strand cDNA. Oligo (dT) was used for reverse transcribing RNA with polyA tail and random primer was used for overall RNA level detection. For the detection of IAV viral vRNA, cRNA and mRNA, uni12 (5’-AGCAAAAGCAGG-3’), uni13 (5’-AGTAGAAACAAGG-3’) and oligo (dT) was used separately for reverse transcription. PrimeSTAR GXL DNA polymerase (R050A, Takara) was used for conventional PCR amplification. *Taq* DNA Polymerase, native (18038042, Thermofisher) was used for amplification of exon-exon junction PCR of *BCAR4*. PCR reactions were prepared as per manufacturer’s instructions. Fast SYBR Green Master Mix (4385612, Thermofisher) was used for qRT-PCR and gene expression was shown in 2^^-Δ^. All primers and oligos sequences used in the study were listed in Table 3.

**Table 3.**
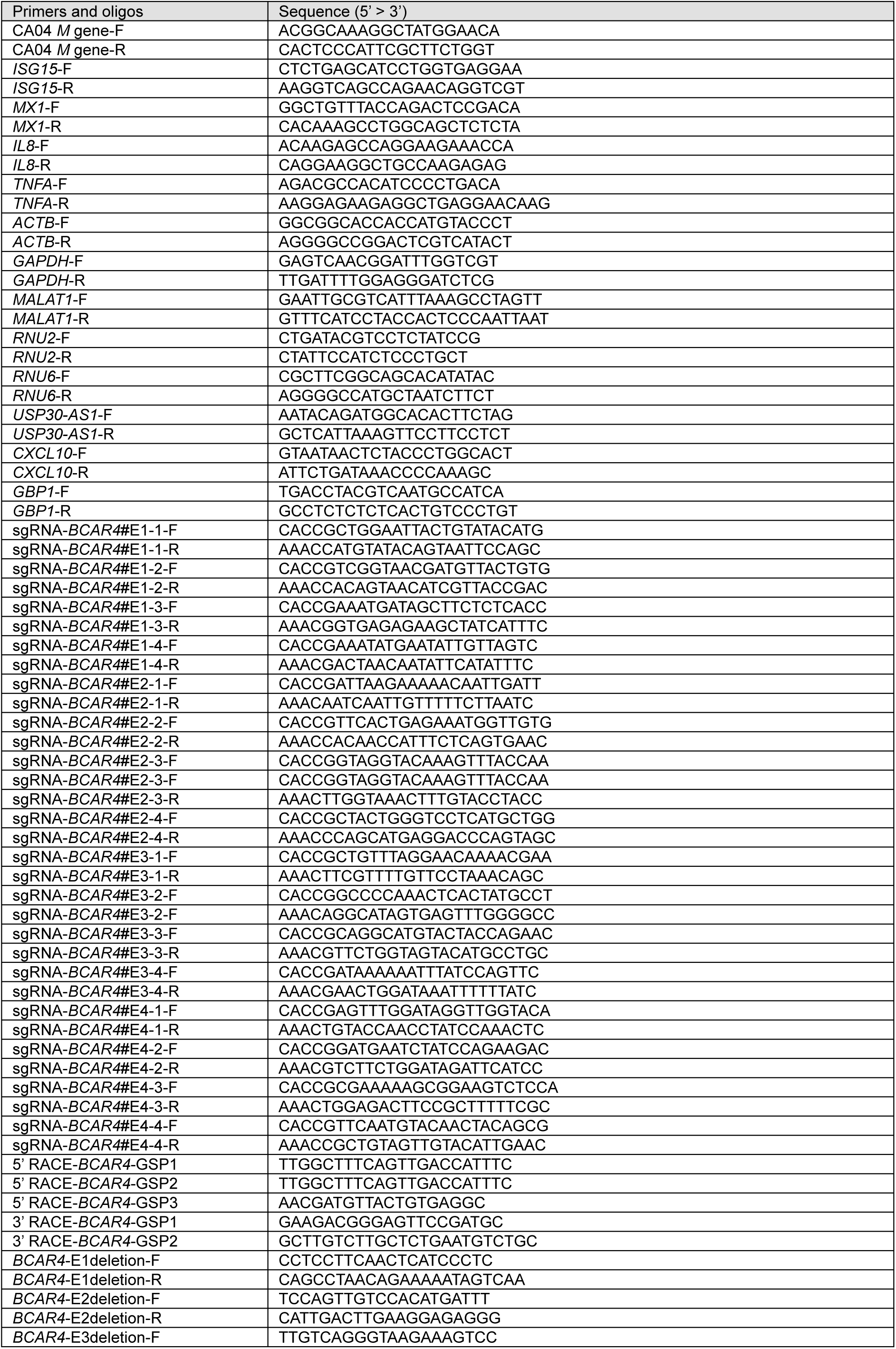
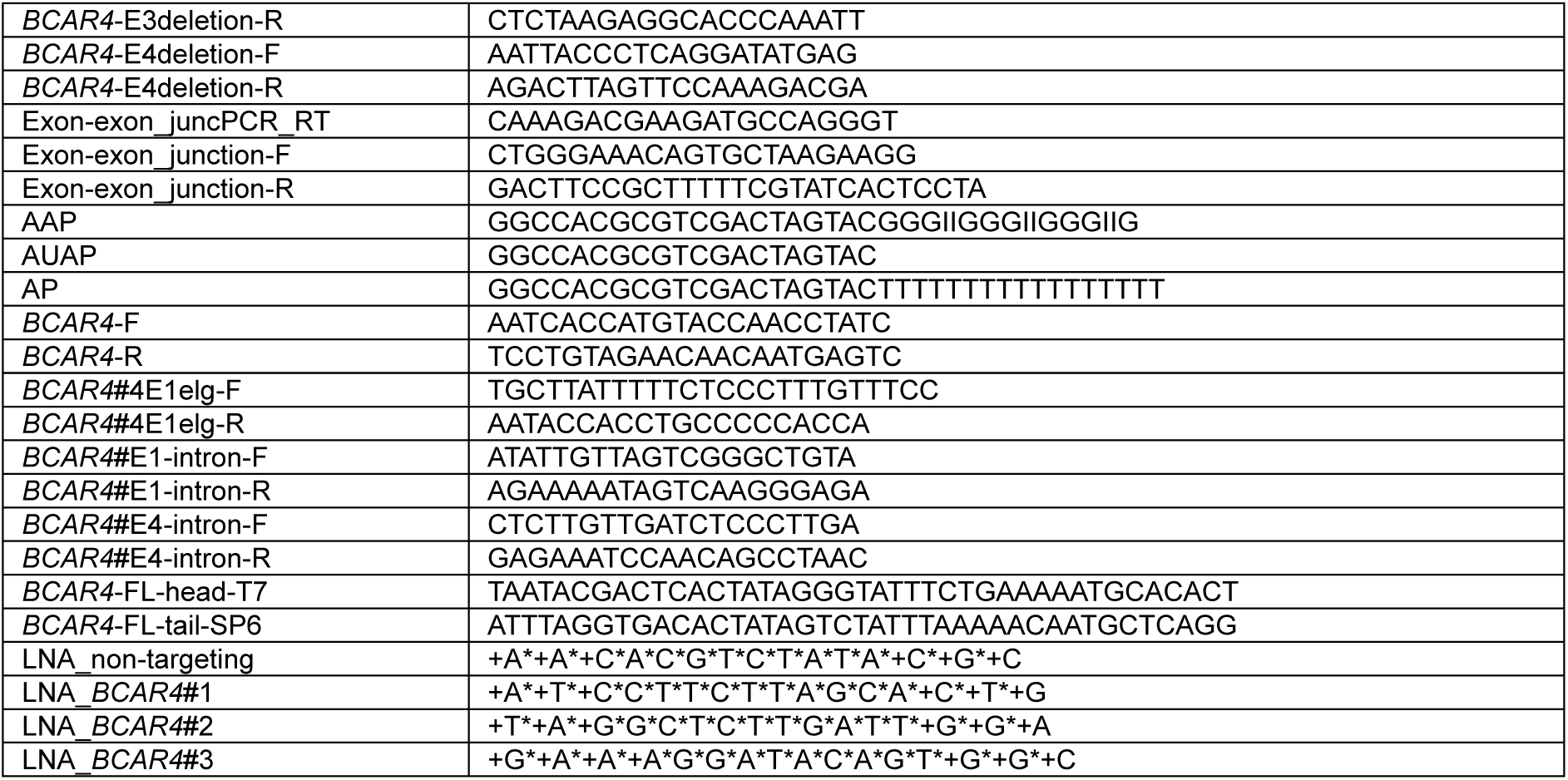
Primers and oligos sequences used in the study.

### 5’/3’ Rapid Amplification of cDNA Ends (RACE) and *BCAR4* exon-exon junction PCR

The 5’/3’ RACE was performed based on the 5’/3’ RACE Systems (18374-058;18373-019, Thermofisher). For 5’ RACE, A549 cells were seeded into T75 flasks with 70% confluence before infection. Next day, cells were either infected with A/California/04/09 (H1N1) of M.O.I. of 1 or treated with PBS as mock infection. The A/California/04/09 (H1N1) and mock infection were both conducted in duplicate. IAV infected or mock infected cells were harvested at 24 hours post infection by using Trizol (15596026, Thermofisher) and RNA was extracted and purified according to Trizol-based protocol. RNA was reversely transcribed into cDNA by using *BCAR4* gene specific primer (GSP). The reverse transcribed cDNAs were capped with dCTP at 5’ end by using terminal deoxynucleotidyl transferase (EP0161, Thermofisher) based on manufacturer’s instructions. First round 5’ RACE PCR amplification was performed by using *BCAR4* GSP together with G-enriched anchor primer (AAP) for 25 cycles. Gel electrophoresis was performed with the first round PCR product. If no target DNA band was exhibited in the gel, a second round PCR was required to perform. For second round PCR, the reaction was prepared by using nested gene specific primer and nested anchor primer (AUAP) with 100-fold diluted first round PCR product as input for 30 cycles of amplification.

For 3’ RACE, cells were seeded in 24-wells plates with 70% confluence. Next day, cells were transfected with a specific LNA or infected with A/California/04/09 (H1N1) infection or treated with PBS-based mock infection, to activate the expression of *BCAR4*. After treatment, cells were harvested, and RNA was isolated and purified by using RNeasy Mini Kit. RNA was next reverse transcribed into cDNA with *BCAR4* GSP in the consensus region of last exon. The first round of 3’ RACE PCR was then performed by using another *BCAR4* GSP with an anchored primer (AP). Gel electrophoresis was performed with the first round PCR product. If no target DNA band was exhibited in the gel, a second round PCR was required to perform, similar to 5’ RACE.

For *BCAR4* exon-exon junction PCR, two experiments were performed separately for Nanopore sequencing or clone library construction. For samples prepared for Nanopore sequencing, RNA from 5’ RACE preparation steps were used and reverse transcribed into cDNA by using GSP in the *BCAR4* last exon consensus region across all transcript variants. Next, two different *BCAR4* GSPs (one was in the consensus region of *BCAR4* first exon, and another was in the consensus region of *BCAR4* last exon) were used for 35 cycle PCR amplification. Gel electrophoresis was conducted with the product of exon-exon junction PCR for visualization. *BCAR4* exon-exon junction PCR for clone library construction was described in a later section. Primers used in the section were listed in Table 3.

### Nanopore sequencing

The PCR product of 5’ RACE and *BCAR4* exon-exon junction PCR were sequenced by using Nanopore long reads sequencing. DNA products from 5’ RACE and *BCAR4* exon-exon junction PCR were subjected to gel purification by using Qiagen Gel Extraction Kit (28706X4, Qiagen). PCR products from 5’ RACE and exon-exon junction PCR from different replicates were barcoded separately and then pooled together into a master mix. Direct cDNA Sequencing Kit (SQK-DCS109, Nanopore), together with Native Barcoding Expansion 1-12 (EXP-NBD104, Nanopore) kit were used to prepare cDNA library according to manufacturer’s instructions. Nanopore long reads sequencing was loaded in Flow Cells and sequenced in MinION sequencer for 72 hours. Base calling was performed in fast5 files by using Guppy (version 5.0.14) (29) and produced fastq files were mapped into human reference genome (GRCh37_latest_genomic.fna) with minimap2 (version 2.22) (30) by using default parameters. Mapped bam files were visualized by using IGV genome browser.

### Clone library construction and Sanger sequencing

To determine dominant RNA transcript of *BCAR4* induced by IAV infection, PCR product of *BCAR4* head-to-tail exon-exon junction PCR from cDNA reversely transcribed from RNA in either A/California/04/09 (H1N1) infected A549 cells or mock infected A549 cells in triplicate was used to construct a clone library. To reduce potential bias introduced by different amounts of cDNA input, we used low (1 μl), medium (3 μl) and high (5 μl) volumes of cDNA input separately for PCR.

PCR products were next purified by using QIAquick Gel Extraction Kit as per manufacturer’s instructions. Purified PCR products generated with different input volumes in each treatment (infection or not) replicate were then pooled together. Pooled PCR products were ligated into TOPO Cloning Vectors (450031, Thermofisher) based on the manufacturer’s instructions. Ligated vectors were transformed into Top 10 E. coli competent cells and spread in LB plates containing 100 μg/ml Kanamycin. Next day, 72 clones were either picked randomly in the plate of the mock infection group or plate of the IAV infection group. Picked clones were amplified and purified by using TIANprep Mini Plasmid Kit as per manufacturer’s instructions. Purified plasmids were sent to Sanger sequencing, which was performed at Centre for PanorOmic Sciences (CPOS), HKU. The number of each sequenced *BCAR4* transcript was counted and the dominant transcript of *BCAR4* was determined in IAV infection or mock infection.

### High-throughput data

Published high-throughput datasets generated from RNA sequencing and microarray were included based on the study design (IAV infected hosts versus mock infected host) and used host (tissues, organoid, or cell lines of human origin) for infection. In total seven different IAV subtypes from six BioProject from the NCBI SRA database were included. Part of the datasets were downloaded from BioProject PRJNA656848 for investigating the expression of *BCAR4* during infection of IAV with *NS1* segment deletion. Raw reads were first mapped into the human reference genome by using STAR aligner (version 2.7.9a) (31) with default parameters and then were counted by using featureCounts (version 2.0.1) (32) with default parameters except for single or pair end mode based on sequencing type. Differential expression analysis was performed by using DEseq2 (version 1.32.9) in R (version 4.1.1).

### Immunoblot

0.176 x 10^6^ A549 cells were seeded into a 24-well plate one day before infection. Different subtypes of IAV or mock infection were added to the cells the next day based on the study purposes. Supernatant was aspirated and cells were lysed in RIPA buffer (NaCl 150 mM, NP-40 1%, DOC 5%, SDS 0.1% and Tris 50 mM with pH 7.4) containing 1X Halt™ Protease Inhibitor Cocktail, EDTA-Free (87785, Thermofisher). Samples were incubated on ice in a shaker for 10 mins. Lysates were then centrifuged at 12000 g at 4 °C for 10 mins to remove cell debris. Protein lysates were quantified by using BCA (23225, Thermofisher) method as per manufacturer’s instructions. Appropriate volumes of 6X protein loading dye were added to the sample lysate to reach 1X. Cell lysates with 1X protein loading dye were boiled at 95 °C for 10 mins. Loading volume for protein electrophoresis was normalized to the sample with lowest protein concentration. SDS-PAGE gel with different percentages was used based on the size of target protein. Nitrocellulose membrane was used for transferring proteins. Protein was blocked by 3% BSA for 30 mins. Primary antibody was added to the membrane and incubated with shaking overnight at 4 °C. Secondary antibody was added to the membrane and incubated at room temperature in the dark for 30 mins. The membrane then was dried in the dark for 1 hour and scanned in Odyssey 9120 Near-infrared Imager (LI-COR).

### RNA pulldown assay

Full length of the *BCAR4* dominant RNA transcript was *in vitro* transcribed by using the dominant transcript DNA template of *BCAR4* (generated from overlapping PCR). For transcription of sense strand of *BCAR4* dominant transcript, HiScribe™ T7 High Yield RNA Synthesis Kit (E2040S, NEB) was used, while HiScribe™ SP6 RNA Synthesis Kit (E2070S, NEB) was used for transcribing the antisense strand. Bio-16-UTP (AM8452, Thermofisher) was used in the reaction with a ratio of 1:3 relative to standard NTP for integrating biotinylated UTP into *BCAR4* RNA transcripts. The *in vitro* transcription was performed based on manufacturer’s instructions. *In-vitro* transcribed RNA was next purified by using RNA Clean Beads (A63987, Beckman) and diluted in RNA structure buffer (10 mM HEPES pH 7.0, 10 mM MgCl2, RNase-free). To prepare *in-vitro* cell lysate, cells were seeded in T75 flasks and cultured to 70% confluence. The cells then were treated with either A/California/04/09 (H1N1) of M.O.I. of 1 or PBS-based mock infection. Duplicate was applied in each treatment. Cells were incubated for 24 hours. After that, infectious medium or PBS-containing medium was aspirated, and cells were washed with PBS once. Next, cells were lysed with 1 ml RIPA buffer containing proteinase inhibitor and incubated at 4 °C with shaking for 5 mins. Sample lysates were centrifuged at 12000 g at 4 °C for 10 mins to remove cell debris. 1 μg biotinylated RNA probes were added to each cell lysate sample and incubated at 4 °C on a rotator overnight. Typically, 100 μl M-280 Streptavidin Dynabeads (60210, Thermofisher) were prepared for each sample. Streptavidin Dynabeads were washed three times by using RIPA buffer containing 1.5 U/μl RNasin and proteinase inhibitor for 5 mins. The wash buffer was removed and Streptavidin Dynabeads were blocked with 10 μg yeast tRNA and 10 mg BSA containing 0.05 U/μl RNasin and proteinase inhibitor at room temperature for 30 mins. Next, blocking buffer was removed and Streptavidin Dynabeads were diluted in 30 μl RIPA buffer containing 1.5 U/μl RNasin and proteinase inhibitor and transferred to each cell lysate sample containing biotinylated RNA probe. Cell lysate containing biotinylated and Streptavidin Dynabeads were incubated in a rotator at room temperature for 30 min. Streptavidin Dynabeads were then collected in a magnet stand and diluted in a 30 μl RIPA buffer. The RIPA buffer containing beads was boiled at 95 °C for 5 mins and beads were then removed through a magnet stand. Sample lysates containing RNA probe enriched proteins were then preserved at -80 °C for further analysis.

### Mass spectrometry

Protein samples from RNA pull-down assay were loaded into lanes of an SDS-PAGE gel consisting of 3% of stacking gel and 8% of resolving gel. Loaded protein samples were electrophoresed along lanes under 60 Am and electrophoresis was stopped immediately after all samples migrated from stacking gel into resolving gel. Next, gel with stacked protein samples was subjected to Coomassie Blue (1610400, BioRad)-based staining and destaining. Destained gel was sent to Proteomics and Metabolomics Core, CPOS, HKU for exercising and mass spectrometry analysis.

### Immune stimulation assay or splicing inhibition assay

Immune stimuli including interferon α2A (H6041-10UG, Sigma), interferon β (IF014, Sigma), interferon γ (GF305, Sigma), LPS (L2630-10MG, Sigma) and conditioned medium were used to stimulate cells for immune response. Conditioned medium was collected from the MEM cell culture medium of IAV infected cells. In brief, a pool of immune or antiviral molecules was secreted into cell culture medium by cells in response to IAV infection. The viruses in the medium were then filtered away by using Amicon Ultra-15 Centrifugal Filter Unit (UFC910024, Sigma). For interferon and LPS stimulation, a gradient of concentrations of stimulus was treated to cells separately for 6 hours or 24 hours. Cells were harvested after treatment for qRT-PCR analysis. For conditioned medium treatment, virus-free conditioned medium was added to the cells and cells were harvested at 0, 2, 4 and 8 hours after treatment. Madrasin (SML1409-5MG) was used for inhibiting cellular spliceosomes. 50 nM Madrasin was treated with cells for 24 hours and treated cells were harvested after for downstream analysis.

### Statistical analysis

Student’s-t test was used for comparing the difference between two independent groups. One-way ANOVA was applied for detecting the difference among three different groups and *LSD*-test was used as a host-hoc test if multiple tests were needed. Two-sided significance level was applied in the study.

## Acknowledgements

We acknowledge the research computing service provided by HPC, HKU. We also thank the contributors that generated published datasets included in the study. This study was supported by the Theme-based Research Scheme of the Research Grants Council, HKSAR Government (T11-712/19-N), the Health and Medical Research Fund, HKSAR Government (CID-HKU2), and the US National Institute of Allergy and Infectious Diseases, National Institutes of Health (HHSN272201400006C), and InnoHK, an initiative of the Innovation and Technology Commission, the Hong Kong Special Administrative Region (Ref: C2i).

